# Polypyrimidine Tract Binding Protein 1 regulates the activation of mouse CD8 T cells

**DOI:** 10.1101/2022.03.03.482829

**Authors:** Vanessa D’Angeli, Elisa Monzón-Casanova, Louise S. Matheson, Özge Gizlenci, Georg Petkau, Clare Gooding, Rebecca V. Berrens, Christopher W. J. Smith, Martin Turner

## Abstract

We show that the RNA-binding protein Polypyrimidine Tract Binding Protein 1 (PTBP1) is dispensable for the development of naïve mouse CD8 T cells, but is necessary for the optimal expansion and production of effector molecules by antigen-specific CD8 T cells *in vivo*. PTBP1 has an essential role in regulating the early events following activation of the naïve CD8 T cell leading to IL-2 and TNF production. It is also required to protect activated CD8 T cells from apoptosis. PTBP1 controls alternative splicing of over 400 genes in naïve CD8 T cells in addition to regulating the abundance of ∼200 mRNAs. PTBP1 is required for the nuclear accumulation of c-Fos, NFATc2 and NFATc3, but not NFATc1. This selective effect on NFAT proteins correlates with PTBP1-promoted expression of the shorter Aβ1 isoform and exon 13 skipped Aβ2 isoform of the catalytic A-subunit of calcineurin phosphatase. These findings reveal a crucial role for PTBP1 in regulating CD8 T cell activation.

## Introduction

Upon antigen recognition through peptide presentation by MHC I and co-stimulation a series of signalling pathways are triggered in CD8 T cells which ultimately lead to cytoskeleton remodelling, metabolic reprogramming, and transcription factor activation [1]. The Calcium, MAP kinase and PKC signalling pathways function by inducing nuclear translocation of the Nuclear Factor of Activated T cells (NFAT), Activator Protein 1 (AP-1) and Nuclear Factor Kappa-light-chain-enhancer of activated B cells (NFκB) transcription factors [2]. These coordinate the expression of multiple genes involved in a programme of proliferation and differentiation. In addition, they induce expression of pro-inflammatory cytokines, including tumour necrosis factor (TNF), interferon gamma (IFNγ), and interleukin-2 (IL-2) [3], [4]. Cytokine expression by CD8 T cells is also regulated post-transcriptionally; this has been shown to be context-dependent and is orchestrated by Calcium, MAPK and PKC pathways, which may coordinate the activities of RNA binding proteins (RBPs) that regulate the stability and translation of mRNA [5].

Polypyrimidine Tract Binding Protein 1 (PTBP1) is a ubiquitously expressed RBP which regulates alternative splicing (AS), polyadenylation, mRNA abundance and internal ribosome entry site-mediated translation [6],[7]. Previous work has shown that *Ptbp1* is important in B lymphocytes for the germinal centre reaction [8],[9] and that it is redundant with its paralogs *Ptbp2* in early B cell development [10] and with *Ptbp3* in mature B cells [11]. PTBP1 also promotes CD154 (CD40-ligand) expression by mouse CD4+ T cells by binding the 3’UTR of *Cd40lg* mRNA [12]. In human CD4 T cells, PTBP1 has been found to be necessary for the optimal signalling via the NFκB and ERK1/2 pathways [13]. Upon knockdown of PTBP1, CD4 T cells showed a reduced proliferation, survival, and expression of effector molecules [13]. PTBP1 also regulated 3’ end formation of the human CD5 transcript in T cells [14]. Whether PTBP1 regulates CD8 T cell responses is unknown.

Here we identify PTBP1 as a regulator of the early signalling events that occur upon CD8 T cell activation required for the production of IL-2 and TNF. In the absence of PTBP1 the accumulation of c-Fos and NFATC3 in nuclei is reduced. We show that PTBP1 regulates the alternative splicing of many genes including the Aβ catalytic subunit of calcineurin, which may contribute to the activation of CD8 T cells.

## Results and Discussion

### PTBP1 is essential for the expansion and differentiation of CD8 T cells *in vivo*

We generated mice with conditional deletion of *Ptbp1* in T cells (*Cd4*^*CreTg*^*Ptbp1*^*fl/fl*^ -abbreviated to P1KO). PTBP1 protein was absent in P1KO CD8 cells from P1KO mice and, as expected, PTBP1-KO CD8 T cells expressed PTBP2 (Supporting information Fig. S1A) due to loss of the repressive effect of PTBP1 on PTBP2 expression [6]. The numbers of naïve CD8 cells in the spleen and lymph nodes of P1KO mice were similar to control *Cd4*^*CreTg*^ mice (Supporting information Fig. S1B). Following introduction of the OT1 transgenic T cell receptor, we found that ∼90% of naïve P1KO OT1 cells lacked PTBP1 protein (Supporting information Fig. S1C). Taken together, these data indicate that the accumulation of naïve CD8 T cells shows no absolute requirement for PTBP1.

To identify cell-intrinsic roles for PTBP1 in CD8 T cell activation, we transferred naïve OT1 cells into recipient mice and subsequently infected them with *Listeria monocytogenes* expressing ovalbumin (attLm-OVA). P1KO OT1 cells showed a severe expansion defect compared to PTBP1-sufficient OT1 cells (Fig. 1A). Upon co-transfer of P1KO and control OT1 cells and infection with attLm-OVA, the expansion of PTBP1-deficient OT1 cells was also greatly reduced (Fig. 1B, C). Transferred P1KO OT1 cells underwent fewer cell divisions than control OT1 cells (Fig. 1D) and more frequently stained positive for active-Caspase-3 (Fig. 1E and Supporting Information S1D) showing that PTBP1 was required for proliferation and protected cells from apoptosis *in vivo*.

**Figure 1.**
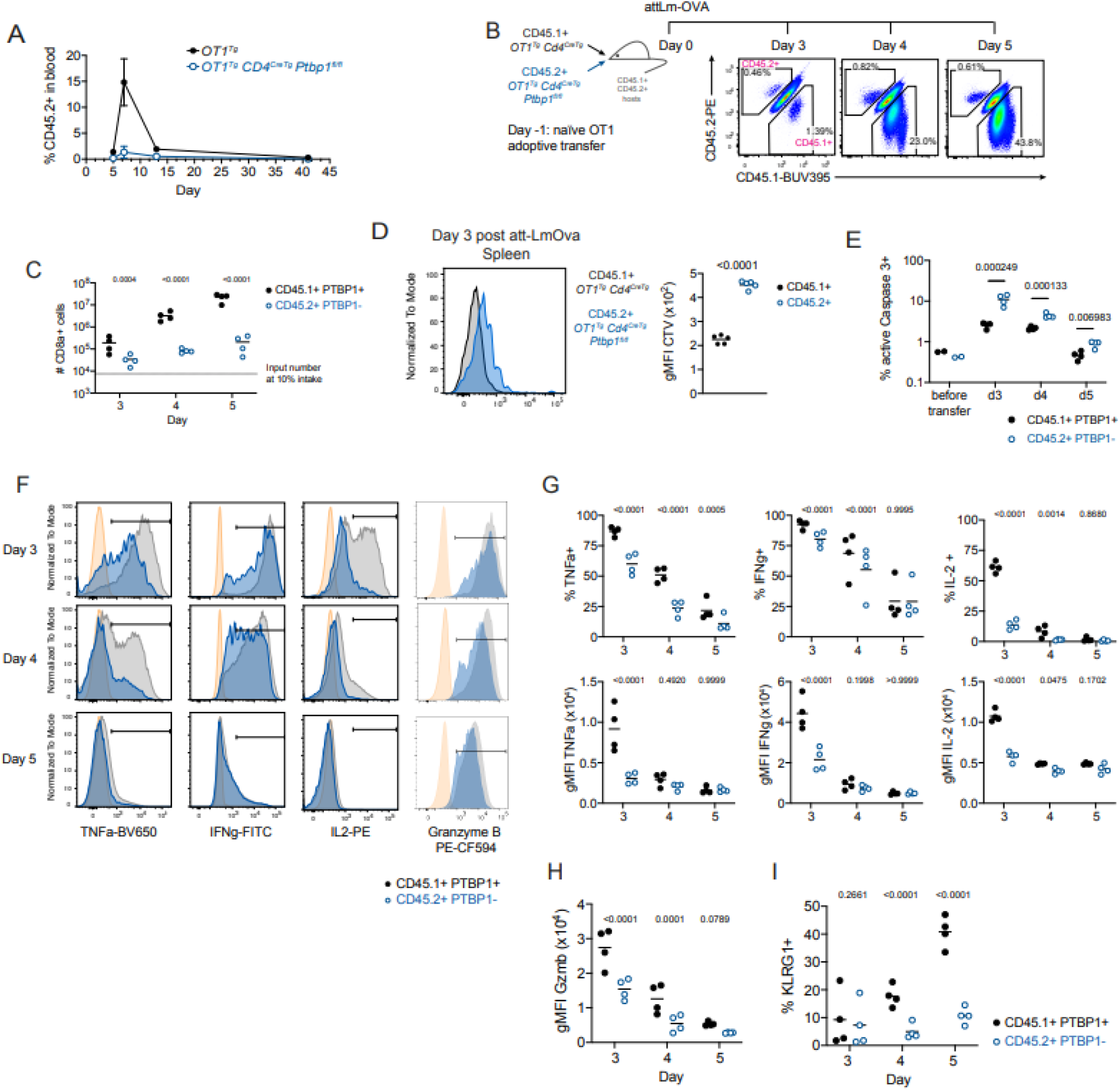
PTBP1 is essential for the expansion and differentiation of CD8 T cells *in vivo*. A) Proportions of CD45.2+ cells (amongst live eFluor780-lymphocytes) identified by flow cytometry in the blood of B6.SJL (CD45.1+) mice which received 1,000 naïve CD8 T cells from OT1 control (CD45.2+ *OT1*^*Tg*^) or OT1 P1KO (CD45.2^+^ *OT1*^*Tg*^ *Cd4*^*CreTg*^ *Ptbp1*^*fl/fl*^) mice and were infected with attLm-OVA one day after OT1 cell transfer. Each symbol shows the mean ± SD of 4 mice per group. Data shown are from one experiment. B) 75,000 naïve CD8 T cells from CD45.1+ *OT1*^*Tg*^ *Cd4*^*CreTg*^ and 75,000 naïve CD8 T cells from CD45.2+ *OT1*^*Tg*^ *Cd4*^*CreTg*^ *Ptbp1*^*fl/fl*^ P1KO mice were adoptively co-transferred into CD45.1+ and CD45.2+ double positive mice which were infected with attLm-OVA one day after cell transfer. Shown are the proportions of CD45.1+ or CD45.2+ cells amongst live (eFluor780-) CD8a+ splenocytes at different days post infection. C) Numbers of CD45.1+ or CD45.2+ cells amongst live (eFluor780-) CD8a+ splenocytes per mouse at different days post infection treated as shown in B. Padj values shown are from Sidak’s multiple correction test post two-way ANOVA from the log-transformed data. D) Cell Trace Violet profiles of CD45.1+ or CD45.2+ CD8 OT1 cells (identified as live eFluor780-and CD8a+) in spleens of CD45.1 and CD45.2 double positive mice which received an adoptive co-transfer of 200,000 naïve CD45.1 control OT1 cells (from *OT1*^*Tg*^ *CD4*^*CreTg*^ mice) and 200,000 naïve CD45.2+ P1KO OT1 cells (from *OT1*^*Tg*^ *CD4*^*CreTg*^ *Ptbp1*^*fl/fl*^ mice) three days post infection with attLm-OVA. Graph shows gMFI of CD45.1+ or CD45.2+ CD8 OT1 cells shown on the left. Each point shows data from one mouse. Lines show arithmetic means. P-value from a two-tailed paired Student’s t-test is shown. Data is from one experiment with 5 CD45.1 and CD45.2 double-positive recipient mice. E) Percentages of active caspase 3+ cells (identified as shown in **Supporting Information S1D**) on different days after att-LmOva infection in mice that received an adoptive co-transfer of control and PTBP1-deficient OT1 naïve CD8 T cells as described in **Figure 1B**. P values shown are from two-tailed paired Student’s t-tests done on the log transformed data. Lines show arithmetic means. Data shown are from one experiment with 4 mice analysed per day. Each point shows data from an individual mouse. F) Cytokine and granzyme B staining of CD45.1+ PTBP1+ (black histograms) or CD45.2+ PTBP1-(blue histograms) live (eFluor780-) CD8+ splenocytes of mice treated as shown in B and re-stimulated with 50 nM SIINFEKL peptide for 3h in the presence of brefeldin A. Orange histograms show PTBP1+ CD8a+ splenocytes from an *OT1*^*Tg*^ mouse. Data shown are from a representative mouse per day. Gates show positive populations. G) Proportions of cytokine expressing cells identified as shown in F and geometric mean fluorescence intensity (gMFI) of the staining within cells identified as positive. Lines show arithmetic means. Data shown are from one experiment with 4 mice analysed per day. Padj values shown are from Sidak’s multiple correction test post two-way ANOVA. H) gMFI of granzyme B staining amongst CD45.1+ PTBP1+ or CD45.2+ PTBP1-live (eFluor780-) CD8+ splenocytes. Lines show arithmetic means. Data shown are from one experiment with 4 mice analysed per day. Padj values shown are from Sidak’s multiple correction test post two-way ANOVA. I) Proportions of KLRG1+ cells amongst CD45.1+ PTBP1+ or CD45.2+ PTBP1-cells in spleens of mice treated as described in B. Gating strategy is shown in **Supporting Information Fig.S1E**. Lines show arithmetic means. Data shown are from one experiment with 4 mice analysed per day. Padj values shown are from Sidak’s multiple correction test post two-way ANOVA.

Amongst the few P1KO cells present both the amounts and the proportions of cells producing TNF, IFNγ and IL-2 were reduced (Fig. 1F and G). Moreover, while all PTBP1-deficient cells contained granzyme B (Fig. 1F) there was a ∼two-fold reduction (Fig. 1H). P1KO OT1 cells present at days four- and five-post-infection also had diminished expression of KLRG1 (Fig. 1I and Supporting Information S1E). Thus, PTBP1 contributes to the differentiation of effector CD8 T cells and is necessary for the production of effector molecules *in vivo*.

### PTBP1 is necessary for optimal activation of CD8 T cells

Following *in vitro* stimulation of naïve P1KO CD8 cells with anti-CD3 and anti-CD28 antibodies the number of cells that have undergone two or more cell divisions were reduced compared to control CD8 cells (Supporting information Fig. S2A). This defect was accompanied by reduced amounts of IL-2 in the culture supernatants (Supporting information Fig. S2B). PTBP1-deficient CD8 T cells responded to the addition of recombinant IL-2 showing increased proliferation (Supporting information Fig. S2C and S2D), but the %-divided cells and the proliferation index were always less than control cells (Supporting information Fig. S2D). The frequencies of active-Caspase-3+ cells were similar between naïve or 24h stimulated P1KO and control CD8 cells (Supporting information Fig. S2E). However, at 48 hours, both non-divided and proliferating PTBP1-deficient CD8 T cells had increased frequencies of active-Caspase-3+ cells at all division numbers compared to controls (Supporting information Fig. S2E). Thus, even when supplemented with IL-2, PTBP1 is necessary for the optimal proliferation and survival of activated CD8 T cells *in vitro*. The reduced proliferation, IL-2 production and increased apoptosis of antibody-stimulated non-transgenic cells reflects the *in vivo* behaviour of the P1KO OT1 cells. The results are consistent with findings in human CD4+ T cells following retroviral delivery of *PTBP1* RNAi and point to a requirement for PTBP1 function long-after stimulation in maintaining cell survival [13]. Moreover, as reported in human CD4+ T cells, we find in mouse CD8+ T cells that PTBP1 is required for the optimal production of IL-2.

### PTBP1 regulates early signalling events in CD8 T cells

To investigate further the role of PTBP1 in the early events following CD8 T cell activation, we measured TCR induced expression of the transcription factor NR4A1. NR4A1 was reduced in PTBP1-deficient CD8 T cells compared to control (Fig. 2A). Additionally, the production of IL-2 and TNF by P1KO OT1 cells was impaired when cells were stimulated with peptide (Fig. 2B). The production of TNF and IL-2 following treatments of P1KO CD8 cells with PMA+ Ionomycin was also defective pointing to a defect in T cell activation that cannot be overcome through bypass of the receptor proximal transduction pathways (Fig. 2B). Previous work in activated human CD4+ T cells, indicated a requirement for PTBP1 for optimal I□B□ degradation [13]. The nuclear translocation of p65-RelA is a sensitive measure of TCR signal transduction [2], yet we found this to be similar in control and P1KO CD8 cells (Fig. 2C). By contrast, the amounts of c-Fos, a component of AP-1, was reduced in PTBP1-deficient CD8 T nuclei (Fig. 2D). The reduced nuclear accumulation of c-Fos might be a consequence of a reduced amount of c-Fos protein. These findings suggest a role for PTBP1 in regulating the early events upon TCR signalling with an impact in the optimal expression of cytokines.

**Figure 2.**
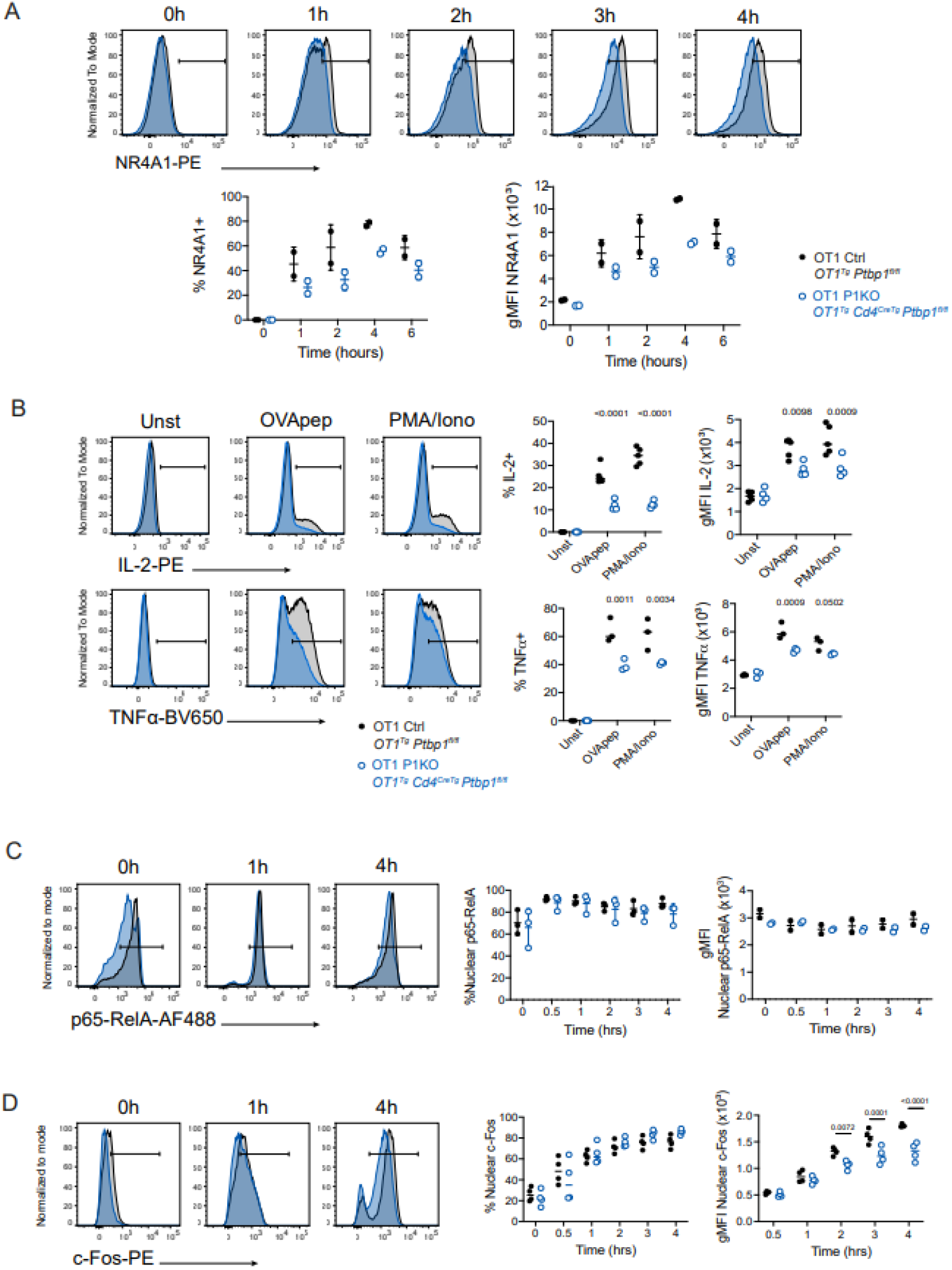
PTBP1 regulates early signalling events in CD8 T cells. A) Histograms showing NR4A1 staining of control (black histograms) or PTBP1-deficient (blue histograms) naïve CD8 T cells activated *ex vivo* with SIINFEKL peptide (10nM) for the time indicated. Gates show positive populations. Below, graphs show proportions of NR4A1-expressing cells and geometric mean fluorescence intensity (gMFI) of the staining within cells identified as positive. Each dot represents a biological replicate. Data shown is from one representative experiment out of three. B) Histograms (left) show IL-2 and TNFα staining in unstimulated (unst) CD8 T cells (eFluor780^-^CD8a^+^TCRb^+^) and stimulated with SIINFEKL peptide (OVApep) or PMA/Ionomycin (PMA/Iono) for 4 hours from control (black histograms) or PTBP1-deficient (blue histograms) in the presence of Brefeldin A. Gates show positive populations. Graphs (right) show proportions of IL-2 or TNFα expressing cells and geometric mean fluorescence intensity (gMFI) of the staining within cells identified as positive. Each dot represents a biological replicate. Data shown is from one representative experiment out of two. C) Histograms (left) show p65-RelA staining of isolated nuclei from control (black histograms) or PTBP1-deficient (blue histograms) naïve CD8 T cells cocultured with bulk splenocytes pulsed with 10 nM of OVA peptide at the time indicated. Gates show positive populations. Graphs (right) show proportions of nuclear translocation of p65-RelA and geometric mean fluorescence intensity (gMFI) of the staining within cells identified as positive. Each dot represents a biological replicate. Data shown is from one representative experiment out of two. OT-I nuclei were identified as CellTrace Violet^hi^ CD3e^low^. D) Histograms (left) show c-Fos staining of isolated nuclei from control (black histograms) or PTBP1-deficient (blue histograms) naïve CD8 T cells cocultured with bulk splenocytes pulsed with 10 nM of OVA peptide at the time indicated. Gates show positive populations. Graphs (right) show proportions of nuclear translocation of c-Fos and geometric mean fluorescence intensity (gMFI) of the staining within cells identified as positive. Each dot represents a biological replicate. Data shown is from one representative experiment out of two. OT-I nuclei were identified as CellTrace Violet^hi^ CD3e^low^.

### PTBP1 controls alternative splicing of Calcineurin Aβ in CD8 T cells

We performed RNA-seq on the transcriptome of naïve CD8 T cells from P1KO and control mice. In the absence of PTBP1 approximately 100 mRNAs were both increased and decreased in abundance (FDR <0.05, Supporting Information Fig.S3A – Source data 1); as this number was calculated with no thresholds on expression values, we conclude PTBP1 has a modest effect on transcript levels in resting CD8+ T cells. We found changes in AS amongst 453 genes (Supporting Information Fig. S3B – Source data 2) indicating a role for PTBP1 in splicing regulation. Amongst these changes, we focussed on the catalytic subunits of calcineurin, a serine-phosphatase with an important role in TCR signalling.

Resting CD8 T cells express all three paralogs of calcineurin A with Calcineurin-Aα (encoded by *Ppp3ca*) and -Aβ (encoded by *Ppp3cb*), expressed at much higher protein copies/cell than -Aγ (encoded by *Ppp3cc*) (Fig. 3A) [15]. In P1KO CD8 T cells no differences were found in the total abundance of the mRNAs encoding the paralogs (Fig. 3B). However, *Ppp3cb* had increased inclusion of exon-13 in PTBP1-deficient compared to control cells (Fig. 3C and Supporting Information Fig. S3C). PTBP1 binds to the introns flanking exon-13 (Fig. 3C) where it is predicted to promote exon skipping. This is supported by the FPKMs estimated for each isoform by cuffnorm, which indicates a decrease in the exon 13-skipped *Ppp3cb-201* isoform in PTBP1-deficient cells (Fig.3D). In addition, *Ppp3cb* has an alternative polyadenylation site in intron-12. This generates an extended exon-12 (transcript *Ppp3cb-206*) encoding a shorter protein termed Aβ1 with an alternaitve C-terminus with distinctive localisation and function and the absence of an inhibitory domain present in the Aβ2 variant [16],[17]. Previous work had indicated this polyadenylation event can be can be regulated by Muscleblind Like Splicing Regulator 1 [18]. PTBP1 binds to the extended exon-12 upstream of the intronic polyadenylation site (Fig. 3C) where it is predicted to promote the use of this alternative polyadenylation site [14]. The FPKM for this isoform is decreased in PTBP1-deficient cells (Fig. 3D). We performed RT-PCR to validate the alternative splicing and polyadenylation of *Ppp3cb* in P1KO naïve CD8 T cells. This confirmed a significant increase in the proportion of transcripts that included exon 13, concomitant with decreased skipping of this exon, and a trend towards decreased usage of the alternative polyadenylation site, although this did not reach significance (Supporting Information Fig. S3D-F). Decreased usage of this alternative polyadenylation site in P1KO cells was further supported by long read Oxford Nanopore Technologies RNA sequencing in naïve and activated CD8 T cells (Supporting Information Fig. S3G-H), albeit many reads were truncated and the number of reads that could be unambiguously assigned to isoforms were low. Therefore PTBP1 promotes the expression of the shorter *Ppp3cb-206* isoform encoding Aβ1 and a variant of Aβ2 which lacks 10 amino acids encoded by exon 13.

**Figure 3.**
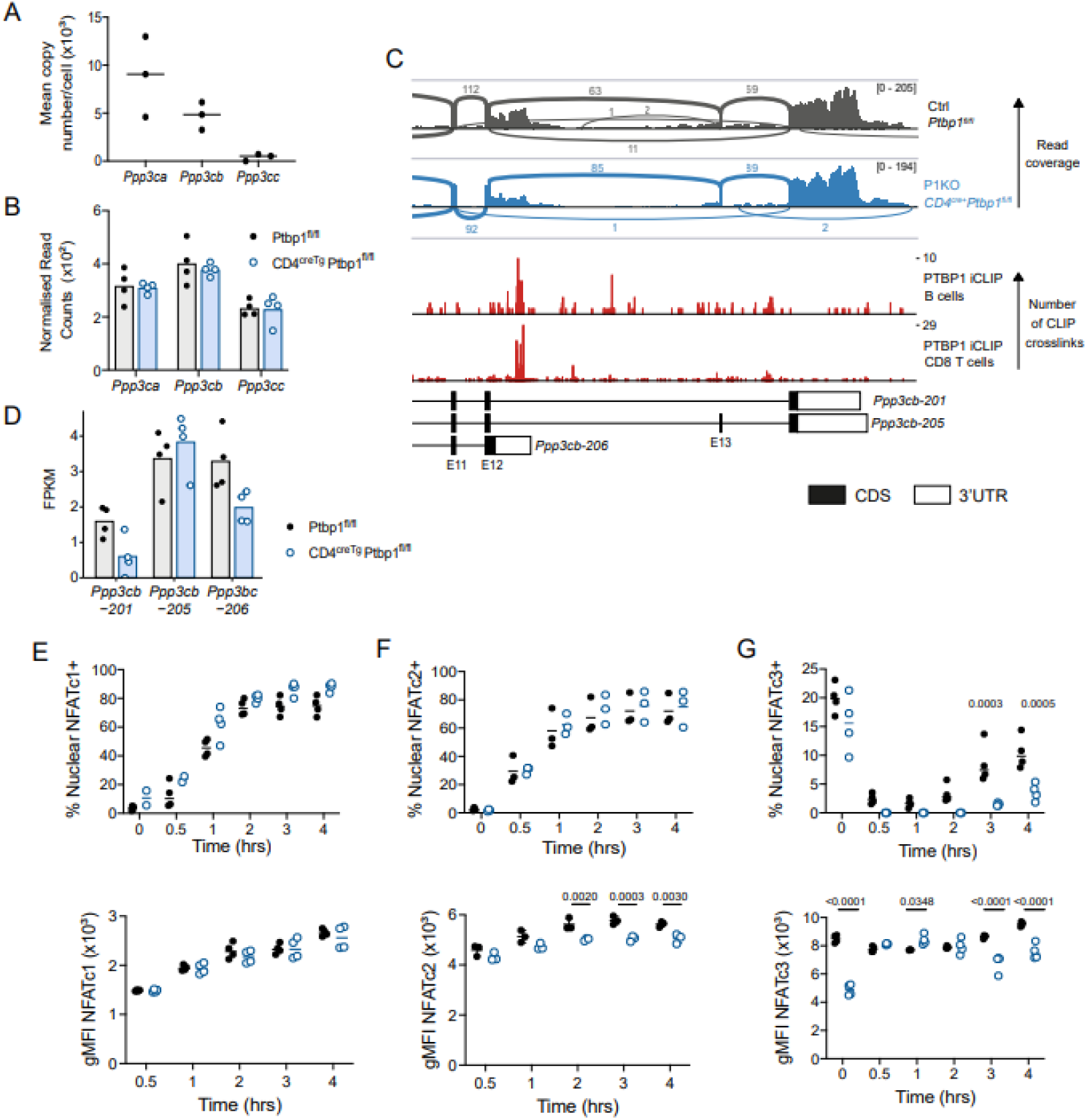
PTBP1 regulated gene expression programs in CD8 T cells. A) Ppp3ca, Ppp3cb and Ppp3cc protein levels analysed by mass spectrometry by Marchingo et al. [15] in naïve CD8 T cells. B) Abundance of *Ppp3ca, Ppp3cb* and *Ppp3cc* mRNA (Normalised Reads Counts from DESeq2 analysis) in naïve CD8 T cells from control (*Ptbp1*^*fl/fl*^) and P1KO mice (*Ptbp1*^*fl/fl*^ *Cd4*^*CreTg*^). C) Sashimi plot of *Ppp3cb* transcript in naïve CD8 T cells from merged replicate datasets for control (*Ptbp1*^*fl/fl*^) and P1KO mice (*Ptbp1*^*fl/fl*^ *Cd4*^*CreTg*^), showing mRNA-seq read coverage and reads that map to exon–exon junctions (arches; numbers indicate reads that map to that junction), and crosslink sites detected by iCLIP for PTBP1 (below). D) Graph showing FPKM of three different isoforms of *Ppp3cb* (shown in the genome reference **Fig. 3C**) in naïve CD8 T cells from control (*Ptbp1*^*fl/fl*^) and P1KO mice (*Ptbp1*^*fl/fl*^ *Cd4*^*CreTg*^). E-G) Graphs showing proportions of nuclear translocation of NFATc1 (D), NFATc2 (E) and NFATc3 (F) and geometric mean fluorescence intensity (gMFI) of the staining within cells identified as positive. OT-I nuclei were identified as CellTrace Violet^hi^ CD3e^low^. Data shown for NFATc1 and NFATc3 are from one experiment with 4 biological replicates out of two. Data shown for NFATc2 are from one experiment out of three (n=3).

Calcineurin acts on many substrates and is essential for the dephosphorylation and nuclear translocation of NFAT. In the mouse C2C12 myogenic cell line the inclusion of *Ppp3cb* exon 13 has been linked specifically to reduced NFATc3 nuclear translocation [19]. In activated CD8 T cells we found that NFATc1 translocation was normal in the absence of PTBP1 (Fig. 3E and Supporting Information Fig. S4A). This suggested that TCR activated calcium signalling was not globally impaired. However, NFATc2 fluorescence intensity in the nucleus of PTBP1-deficient CD8 T cells was slightly reduced compared to control CD8 T cells indicating reduced translocation (Fig. 3F and Supporting Information Fig. S4B). For NFATc3-staining, stimulation resulted in the rapid loss of nuclear staining and a gradual increase in the frequency NFATc3 between two and four hours that was diminished in nuclei from PTBP1-deficient cells (Fig. 3G and Supporting Information Fig. S4C). As there were no differences in the NFATc1, NFATc2 and NFATc3 transcript isoforms [20]–[22] arising from alternative splicing and polyadenylation in PTBP1-deficient CD8 T cells compared to control cells (data not shown) our data implicate PTBP1 in controlling the selective nuclear translocation of NFAT family members indirectly, via an intermediate. We highlight the control of alternative isoforms of Calcineurin Aβ as a candidate mechanism to mediate this effect.

### Concluding remarks

Our data from CD8 cells is broadly consistent with the findings in human CD4 T cells and indicate an important role of PTBP1 in the early activation events following TCR stimulation. These events require the nuclear translocation of a collection of transcription factors that together stimulate the gene expression programme of the activated T cell [23],[24]. It is likely that PTBP1 will regulate gene expression at times late after activation and this may include genes that protect activated T cells from apoptosis. In the present work we validate *Ppp3cb* as a direct target of PTBP1 and its isoforms as candidates for selective regulators of early events in T cell activation; we anticipate that further regulation mediated by PTBP1 of other transcripts will contribute to T cell activation via alternative splicing, polyadenylation and control of their abundance.

## Materials and Methods

### Mice

Mice were maintained in the Babraham Institute Biological Support Unit. No primary pathogens or additional agents listed in the FELASA recommendations have been confirmed during health monitoring since 2009. Ambient temperature was ∼19–21°C and relative humidity 52%. Lighting was provided on a 12hr light: 12hr dark cycle including 15 min ‘dawn’ and ‘dusk’ periods of subdued lighting. After weaning, mice were transferred to individually ventilated cages with 1–5 mice per cage. Mice were fed CRM (P) VP diet (Special Diet Services) *ad libitum* and received seeds (e.g., sunflower, millet) at the time of cage-cleaning as part of their environmental enrichment. All mouse experimentation was approved by the Babraham Institute Animal Welfare and Ethical Review Body. Animal husbandry and experimentation complied with existing European Union and United Kingdom Home Office legislation.

Mice were derived from crossing the following transgenic strains: *Ptbp1*^*fl/fl*^ (Ptbp1^tm1Msol^) [25], Cd4^CreTg^ (Tg(Cd4-cre)1Cwi) [26], and *OT1*^*Tg*^ (Tg(TcraTcrb)1100Mjb) [27]. All mice used were between 7- and 20-weeks old and were age- and sex-matched within experiments.

### Cell and nuclei isolation

Single cell suspensions were prepared from spleen and lymph nodes by passing the organs through 70 and 40 µm cell strainers consecutively. Naïve CD8 T cell isolation was achieved by depletion of cells bound by biotinylated antibodies: CD4 (GK1.5), CD11b (M1/70), CD11c (N418), CD19 (1D3), B220 (A3-6B3), CD105 (MJ7/18), TER-119, gamma/delta TCR (GL3), Gr1 (RB6-8C5), NK1.1 (PK136), F4/80 (BM8), CD44 (1M7) and magnetic Dynabeads™ M-280 Streptavidin (Thermofisher cat#:11206D).

### Naïve CD8 OT1 cell transfer and att-Lm-OVA infection

For analysis of naïve OT1 cell expansion in the blood (Fig. 1A) 1,000 naïve OT1 cells isolated by fluorescence activated cell sorting (eFluor780^-^, CD8a (53-6.7)-APC^+^, CD45.2(104)-PE^+^, CD62L(MEL-14)-BV421^+^, CD44(IM7)-BB515^-^) and mixed with 10^6^ carrier splenocytes from B6.SJL mice were adoptively transferred into B6.SJL recipient mice. One day later, mice were infected with 5×10^6^ C.F.U. attLm-OVA lacking actA [28] intravenously. For analysis of naïve OT1 cell expansion in the spleens of CD45.1/CD45.2 recipient mice (F1 of B6. SJLxC57BL/6 mice) analysed at days 3, 4 and 5 post infection with 5×10^6^ C.F.U. attLm-OVA (Figure 2B), naïve OT1 cells were isolated by magnetic negative depletion as described above. The numbers of naïve OT1 transferred before attLm-OVA infection are specified in the figure legends.

### *Ex vivo* T cell stimulation

10^5^ Naïve CD8 T cells isolated by magnetic negative depletion as described above were cultured in RPMI 1640 Medium (Dutch modification) with 20 mM HEPES (ThermoFisher cat#:22409-015), 50 µM β-Mercaptoethanol, 100 units/ml penicillin and streptomycin, GlutaMAX and 10% FCS on 96 flat-bottom well plates precoated with anti-CD3 (2C11, BioXcell) and CD28 (37.51, BioXcell) antibodies. For tracking cell division naïve CD8 T cells were labelled with either Cell-Trace Violet (ThermoFisher, cat#:C34557) or Cell-Trace Yellow (ThermoFisher, cat#:C34567), following the manufacturer’s recommendations but labelling cells for only 10 minutes. 20 ng/ml (20 units in 200 µl) recombinant murine IL-2 (PeproTECH cat#:212-12-100UG) was added to cultures when indicated. For measuring cytokine production, splenocytes from OT-I transgenic mice were stimulated with SIINFEKL peptide (N4) (ProImmune, Think Peptides) at the concentration of 10^−7^M or with PMA and Ionomycin. For nuclear staining naïve OT-I transgenic CD8 T cells were labelled with CellTrace Violet and stimulated by SIINFEKL loaded splenocytes in a 96 u-bottom plate. Nuclei were isolated using previously described methods [29].

IL-2 amount from supernatant of naïve CD8 T cells activated for 24 hours with with 5 µg/ml anti-CD3 with 1 µg/ml anti-CD28 antibodies was analysed by Meso Scale Discovery (MSD) assay at the Core Biochemical Assay Laboratory (Cambridge).

### Flow cytometry

Single cell suspensions from the spleen and lymph nodes were prepared by passing the tissues through a 70 µm subsequently a 40 µm cell strainers. Antibody binding to Fc receptors was blocked with the anti-FcγIII 2.4G2 monoclonal antibody. Cell surface staining and the Fixable Viability Dye eFluor780 (eBioscience, cat#: 65-0865-18) was carried out at 4°C. The antibody used were: CD45.2-BV785 (104, BioLegend, cat # 109839), CD45.2-PE (104, eBioscience, cat # 12-0454-82), CD45.1-BUV395 (A20, BD, cat # 565212), CD8a-BUV395 (53-6.7, BD, cat # 563786), CD8a.BUV737 (53-6.7, BD, cat # 564297).

For intracellular staining cells were fixed with BD BD Cytofix/Cytoperm (BD cat#: 554722) and either washed with BD Perm/Wash buffer (BD cat#: 554723) or with PBS, frozen at -80°C in 10%DMSO and 90% FCS at least overnight, washed with PBS, re-fixed with BD BD Cytofix/Cytoperm for 5 minutes on ice and washed with BD Perm/Wash buffer. Permeabilised cells were incubated for 20 minutes at room temperature with anti-FcgIII 2.4G2 antibody in BD Perm/Wash buffer. Intracellular staining was carried out with the following antibodies diluted in BD Perm/Wash buffer: Active-caspase-3-GranzymeB-PE (C92-605, BD, cat #561011), Active-caspase-3-BV650 (C92-605, BD, cat #564096), GranzymeB-PE-CF594 (GB11, BD, cat #562462), IFNg-FITC (XMG1.2, BioLegend, cat #505806), IL2-PE (JES6-5H4, BioLegend, cat #503808), KLRG1-BV421 (2F1, BD, cat #562897), NR4A1-PE (12.14, eBioscience, cat #12-5965), PTBP1-AF647 (clone 1, ThermoFisher, cat #32-4800), PTBP2-AF488 (S43, from Michele Solimena), TNFa-BV650 (MP6-XT22, BioLegend, cat #506333). PTBPs anti-bodies were conjugated to different fluorophores with anti-body labeling kits from ThermoFisher (AF488, cat #A20181; AF647, cat #A20186). Intracellular staining for PTBPs was carried out at least for 4 hours at room temperature or overnight at 4°C. Cytokine staining of splenocytes after attLm-OVA infection was carried out after re-stimulation of CD8 T cell enriched splenocytes with 50 nM SIINFEKL peptide for 3 hours with Brefeldin A (Biolegend, cat#:420601).

Staining of nuclei was carried out with the following antibodies diluted in 1x PBS and 2% FCS and 0.3% Triton X-100 overnight at 4°C: anti-cFos (9F6)-PE (Cell-Signalling), anti-p65-RelA (D14E12)- PE (Cell-Signalling), anti-NFATc1 (7A6)-AF488 (Biolegend), anti-NFATc2 (D43B1)-AF488 (Cell-Signalling), anti-NFATc3 (Cell-Signalling).

### mRNASeq libraries

RNA was extracted from naïve CD8 T cells using the RNeasy Micro Kit (cat# 74004, Qiagen). Naïve T cells were isolated from four female control (*Ptbp1*^*fl/fl*^) and four P1KO (*Cd4*^*CreTg*^*Ptbp1*^*fl/fl*^) mice as described in the CD8 T cell isolation section. 10 ng RNA were used and 7 PCR cycles were carried out to synthesise cDNA with the SMART-Seq® v4 Ultra® Low Input RNA Kit (cat # 634891, Takara). RNAseq libraries were prepared with the Nextera XT DNA Library Preparation Kit (cat# FC-131–1096, Illumina) and carrying out 8 PCR cycles. mRNAseq libraries were sequenced on an Illumina HiSeq2500 on a 2×125 bp paired-end run.

### RNA sequencing

mRNA-seq libraries were run across two lanes of a HiSeq 2500 RapidRun. After demultiplexing, data from the two lanes were merged for each sample before trimming TrimGalore (v0.6.1; https://www.bioinformatics.babraham.ac.uk/projects/trim_galore/), and quality control FastQC (https://www.bioinformatics.babraham.ac.uk/projects/fastqc/) and FastQ Screen (https://www.bioinformatics.babraham.ac.uk/projects/fastq_screen/. Reads were mapped to the GRCm38 using HISAT2 (v2.1.0) [30], suppressing unpaired or discordant alignments and taking into account known splice sites from the Ensembl Mus_musculus GRCm38.90 annotation. FPKM were calculated using cuffnorm from Cufflinks (v2.2.1) [31], with geometric normalisation, using the Ensembl Mus_musculus. GRCm38.96 annotation, and genes with average FPKM < 1 or that encode immunoglobulin or T cell receptor genes were excluded from subsequent analysis

### Differential mRNA abundance analysis

Raw read counts were generated using Seqmonk (v1.45.0; https://www.bioinformatics.babraham.ac.uk/projects/seqmonk/) over mRNA features with merged isoforms, using the Ensembl Mus_musculus GRCm38.96 annotation, assuming unstranded, paired reads. Downstream analysis was performed using R v3.5.3. Genes encoding immunoglobulin or T cell receptor gene segments were excluded from analysis; genes with average FPKM < 1 in both conditions were also excluded. DESeq2 analysis (v1.20.0) [32] comparing naïve control and *Ptbp1* KO was performed using default parameters, with ‘normal’ log2 fold change shrinkage. Significantly differentially expressed genes were defined as those with FDR-adjusted p value < 0.05 and mean normalised read counts > 5.

### Transcriptome-wide Alternative splicing analysis

Fastq files that had been trimmed using Trim Galore as above were further trimmed to 123 bases, discarding any reads below this length and any resulting unpaired reads, using Trimmomatic [33]. The resulting reads were mapped with HISAT2 as above, but only retaining uniquely mapped reads in the BAM files (samtools view - bS -F 4 -F 8 -F 256 -q 20). BAM files were sorted and indexed (samtools v1.9) before running rMATS (v4.0.2, turbo) to compare control and PTBP1 cKO cells, specifying a read length of 123 and that libraries are paired end, using the Ensembl Mus_musculus.GRCm38.96 annotation. rMATS results that only counted reads spanning junctions were used for subsequent analysis. Significantly altered splicing events that were significantly altered for each comparison were defined as those with FDR < 0.05, absolute inclusion level difference > 0.1, and supported by a total of at least 35 reads mapping to the included or skipped form in control or P1KO (sum of four biological replicates), and that are from genes with FPKM > 1 in at least one condition (mean of four biological replicates). Duplicated events where the coordinates of the alternatively spliced exon were identical were filtered to retain only the event with the lowest FDR or, where tied, with the greatest number of supporting reads. Sashimi plots to visualise read coverage and junction-spanning reads were generated using the IGV genome browser, after merging replicate datasets into a single BAM file using samtools.

### PTBP1 iCLIP library preparation and sequencing

CD8 T cells isolated by negative magnetic depletion as described above but omitting the anti-CD44 biotinylated antibody from control (*Ptbp1*^*fl/fl*^) or P1KO mice were stimulated ex vivo with plate-bound anti-CD3 (2C11, 5 µg/ml), anti-CD28 (37.51, 1 µg/ml) and 20 ng/ml murine IL-2 for 24 hours. 9×10^5^ CD8 T cells were stimulated per well in 48-well plates. After 24 hours cells were irradiated 1x with 254 nm UV light (150 mJ/cm2). iCLIP libraries were carried out following an improved iCLIP protocol [34]. Cells were lysed with 50 mM Tris-HCl, pH 7.4, 100 mM NaCl, 1% Igepal CA-630 (Sigma I8896), 0.1% SDS and 0.5% sodium deoxycholate lysis buffer. After sonication between 0.5 mg – 0.2 mg protein lysates in 1 ml were incubated with low amounts of RNAse I (0.05 – 0.01 units, ThermoFisher cat#: EN0602) and 4 units Turbo DNAse (Ambion cat#:AM2238) for 3 minutes at 37 °C. PTBP1-RNA complexes were precipitated with 2 µg of anti-PTBP1 antibody (ThermoFisher clone 1 cat#:32-4800) and 20 µl of Protein A/G beads (Pierce cat#:88802) overnight rotating at 4°C. RNA isolation and cDNA library preparation was carried out as described in Lee et al. [34]. using the L3-IR-App adapter to visualise and retrotranscribe the RNA crosslinked to PTBP1. We carried out four biological replicates from control (*Ptbp1*^*fl/fl*^) and two biological replicates from P1KO activated CD8 T cells.

### iCLIP computational processing

Data from CD8 T cells, and from mouse B cells [8] were processed on the iMaps platform (https://imaps.genialis.com/iclip). Reads were deduplicated based on their random barcodes, trimmed using Cutadapt (v2.4) [35] and mapped to the GRCm38 mouse genome build using STAR (v2.7.0f) [36]. Significant crosslink sites and clusters were identified for each replicate separately, except for mouse B cells where replicates were merged, using the iCount pipeline (v2.0.1dev; https://icount.readthedocs.io/en/latest/index.html). The gene and feature type to which each crosslink site or cluster was assigned were determined by considering all transcript isoforms from the Ensembl Mus_musculus. GRCm38.96 annotation. If multiple genes or feature types were overlapped, the most likely was chosen in a hierarchical way: first, protein coding isoforms were prioritised other transcript biotypes; second, prioritising high confidence isoforms with transcript support level < 4; third, by a hierarchy of feature types (3’UTR > CDS > 5’UTR > intron > non-coding exon > non-coding intron); fourth, higher confidence isoforms (based first on transcript support level and second on whether they have a CCDS) were prioritised. iCLIP data were visualised using a shiny app (https://github.com/LouiseMatheson/iCLIP_visualisation_shiny); all crosslinks merged across replicates are displayed, without filtering for significant sites.

### Long read RNA sequencing library preparation

Libraries were generated essentially according to the CELLO-seq protocol [37] with the following modifications: The addition of UMIs by splint ligation was omitted. Reverse transcription was performed using 1 ng total RNA isolated from a pool of cells. PCR reactions were performed using standard Oxford Nanopore Technologies barcoded primers under the following conditions: 90 °C for 45 s, 20 cycles (90 °C for 15 s, 64 °C for 30 s, 72 °C for 10 min), 72 °C for 5 min. PCR products were purified using 0.6X AMPure XP beads (Beckman).

### MinION sequencing of long read cDNA libraries

Equal quantities of each cDNA sequencing library were pooled using a total of 200 fmol, before sequencing on a MinION R9.4.1 flow cell using the SQK-LSK109 kit (1D sequencing). The standard MinKNOW procedure was used for a 72 hour run, according to manufacturers’ instructions.

### Long read data processing and analysis

Raw reads were basecalled using Guppy (4.0.11), in fast mode. Sequence reads with a mean sequence quality of 7 were classified as passed reads; these were first demultiplexed with the Guppy debarcoder option and then processed with pychopper (2.5.0) in order to identify, orient and trim the full-length cDNA reads. Next, the pychopped reads were aligned with minimap2 (2.17) with splice aware setting (-ax splice), which is compatible with pipelines for further isoform identification analysis. The sequences were aligned to mouse genome reference GRCm38. BAM files for replicate datasets were merged using samtools, before filtering for reads overlapping at least one exon of Ppp3cb, and that contained at least one splice junction. These were visualised and a sashimi plot generated in IGV, and the number of reads that could be unambiguously assigned to isoforms including exon 13, skipping exon 13, or utilising the alternative 3’UTR and polyadenylation site within the extended exon 12 were counted manually.

### RT-PCR

cDNA using 0.5 µg total RNA, 50 ng of oligo(dT), 250ng of random hexamers and 5 units of Transcriptor Reverse Transcriptase (Roche) was prepared as described in manufacturer’s protocol. PCRs with 50 ng of prepared cDNAs were carried out using Jumpstart Taq polymerase (Sigma) according to manufacturer’s protocol. Primers binding to exons 11 (forward; 5-TGCAGCCCGGAAAGAAATCA), 14 (reverse; 5-GTATGTGCGGTGTTCAGGGA) and to the alternative 3’UTR in the extended exon 12 (reverse; 5-GAGCTGCTCCCCATATCACC) of *Ppp3cb* were included in each reaction. Cycling parameters were 94°C for 30 seconds, 60°C for 30 seconds and 72°C for 60 seconds for 30 cycles. Minus RT cDNA for each sample were included. For visualisation and quantification of the PSI values, PCR products were resolved in a QIAxcel Advanced System (QIAGEN) and PSI calculated within the QIAxcel ScreenGel software.

## Acknowledgements

We thank Michele Solimena for *Ptbp1* mice and anti-PTBP2 antibodies; Martha Cyert and Idil Ulengin-Talkish for helpful discussions; the Babraham Institute Biological Support Unit, Sequencing, Flow Cytometry and Bioinformatics Facilities for assistance, the Finkelstein Lab (https://github.com/finkelsteinlab) for the BioRxiv template and Sarah Collison for formatting the manuscript. This study was supported by funding from the Biotechnology and Biological Sciences Research Council (BBSRC) (BB/P01898X/1; BBS/E/B/000C0407; BBS/E/B/000C0428; the BBSRC Core Capability Grant to the Babraham Institute; and a Wellcome Investigator award (200823/Z/16/Z) to M.T. V. D’A was supported by a Cambridge Commonwealth, European and International Trust studentship. R.V.B. was supported by a Sir Henry Wellcome Fellowship (no. 213612).

## Author Contributions

E.M.C., C.W.J.S. and M.T. conceptualised the study; E.M.C, G.P., O.G., C.G. and V. D’A carried out experiments; E.M.C., V. D’A, O.G. and L.S.M. analysed data; R.V.B. and O.G. developed methodology; E.M.C., C.W.J.S. and M.T. acquired funding for the study; V. D’A, E.M.C and M.T. wrote the manuscript with input from all the authors.

## Conflicts of Interest

The authors declare no commercial or financial conflicts of interest.

## Data availability statement

The data that support the findings of this study are openly available under the GEO accession GSE190512.

## Notes

### Competing Interest Statement

The authors have declared no competing interest.

