## supplementary figures for "Polypyrimidine Tract Binding Protein 1 regulates the activation of mouse CD8 T cells"

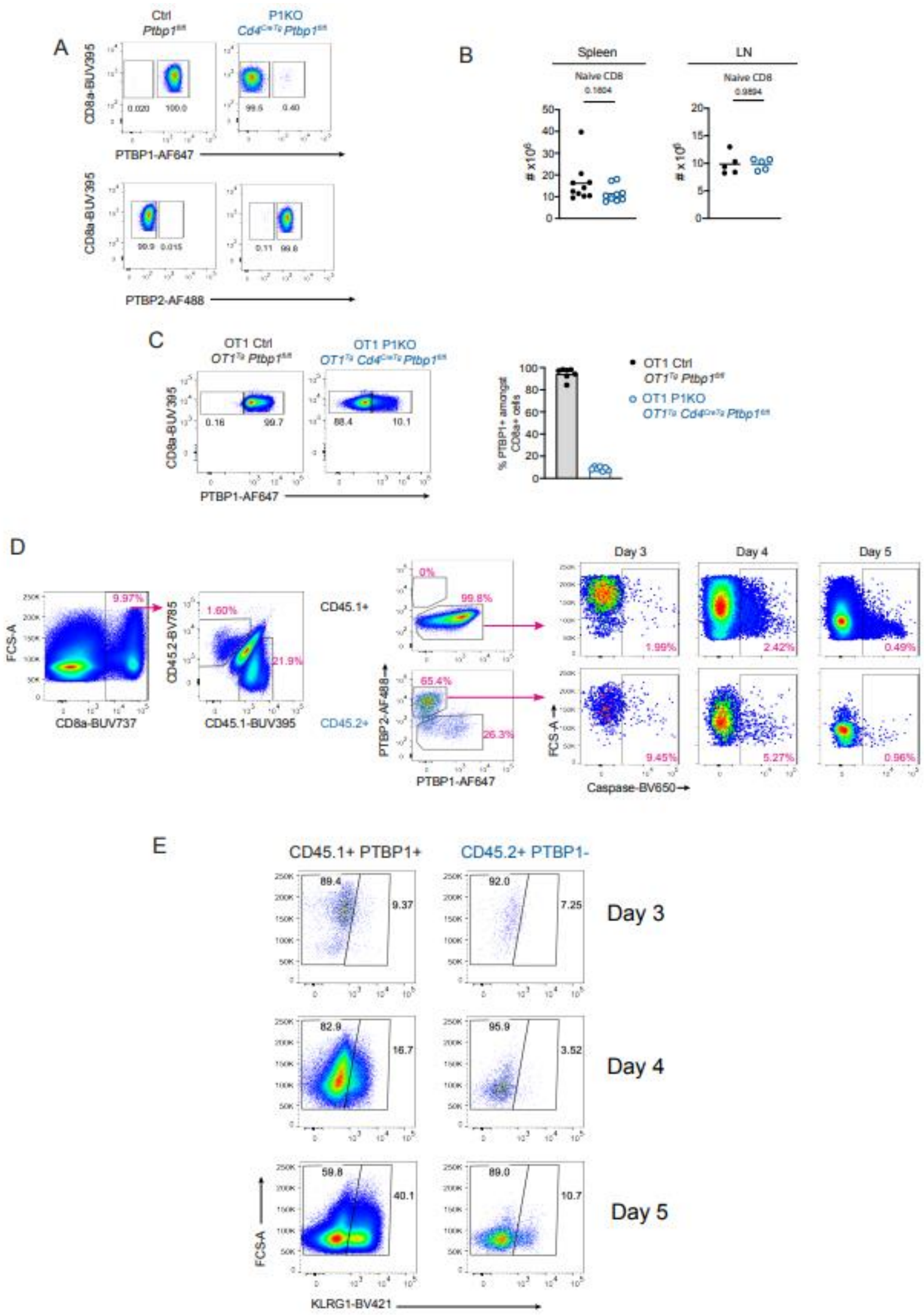


**Supporting Information Figure S1 - PTBP1 is essential for the expansion and differentiation of CD8 T cells *in vivo*** A) Flow cytometry staining of PTBP1 and PTBP2 in splenic CD8 T cells (eFluor780$^-$, TCRb$^+$, CD8a$^+$) from control mice (Ctrl, *Ptbp1*$^{fl/fl}$) and P1KO mice (*Ptbp1*$^{fl/fl}$ *Cd4*$^{CreTg}$). Data shown are from a representative mouse per genotype of 10 mice analysed. B) Numbers of naïve CD8 (eFluor780$^-$, TCRb$^+$, CD8a$^+$, CD62L$^{high}$) T cells in the spleen and peripheral (axial, brachial and inguinal) lymph nodes assessed by flow cytometry. Data shown are from two (spleen) and one (lymph nodes) independent experiments. Each symbol shows data from one mouse. Lines show arithmetic means. Padj values shown are calculated with Tukey's multiple comparison test carried out after one-way ANOVA. C) Flow Cytometry and proportions of OT1 naïve CD8 (eFluor780$^-$, TCRVb5.1, 5.2$^+$, CD8a$^+$, CD62L$^{high}$ and CD44$^{low}$) T cells stained with anti-PTBP1 antibody from control mice (OT1 Ctrl, *OT1*$^{tg}$ *Ptbp*$^{1fl/fl}$) and P1KO mice (*OT1*$^{tg}$ *Ptbp1*$^{fl/fl}$ *Cd4*$^{CreTg}$). Data shown are from a representative mouse per genotype of 7 mice analysed in two independent experiments. Each point shows data from one mouse. D) Gating strategy of active caspase 3+ cells. Shown are splenocytes at day 4 post Lm-Ova infection from mice that received an adoptive co-transfer of control and PTBP1-deficient OT1 naïve CD8 T cells as described in **Figure 1B**. E) Gating strategy of KLRG1+ cells amongst live (eFluor780-) CD8a+ splenocytes from mice treated as described in **Figure 1B** that are either CD45.1+ and PTBP1+ or CD45.2+ and PTBP1-. Data from a representative mouse per day post infection with attLm-OVA are shown. Numbers show percentages of gated cells.

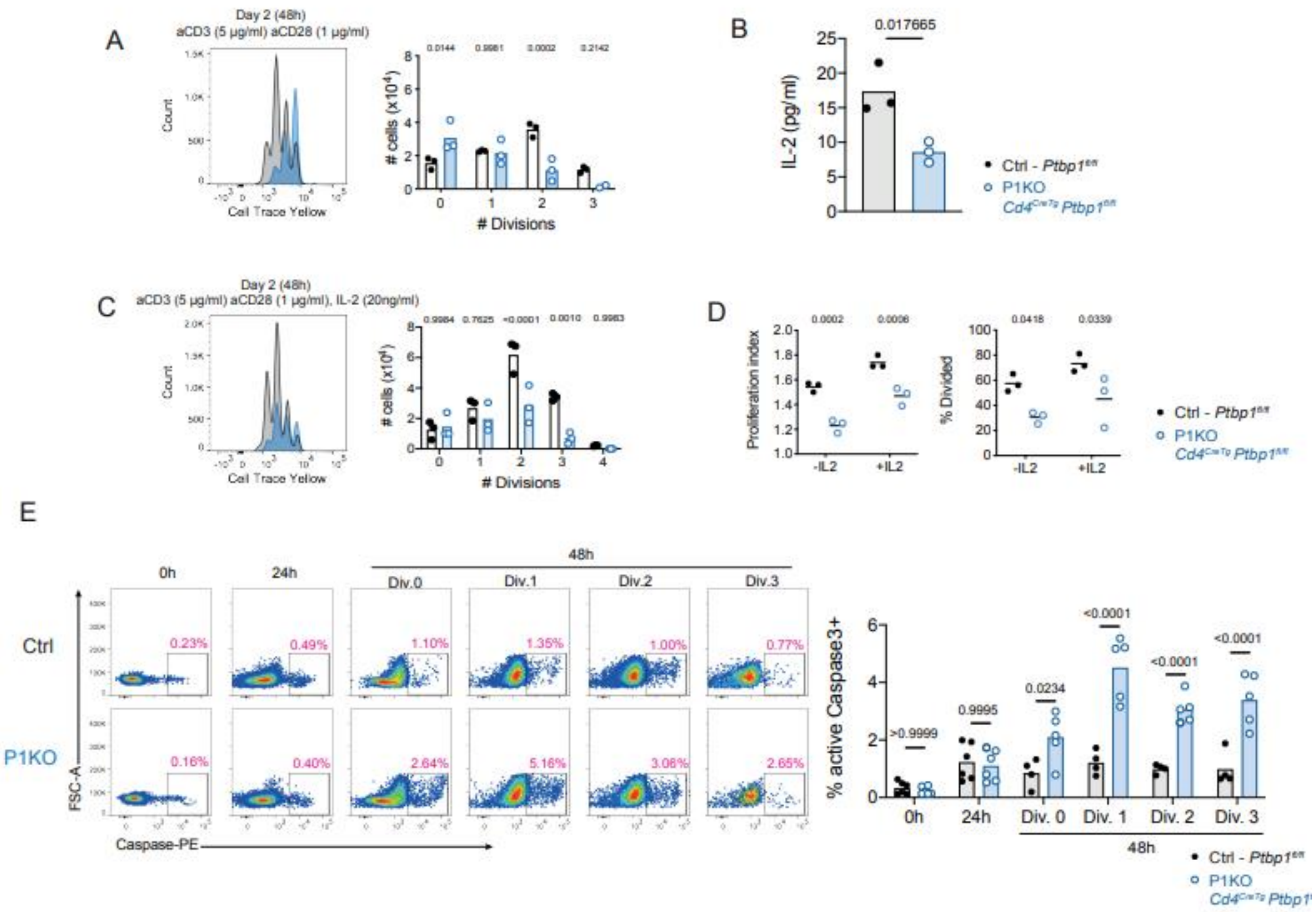


**Supporting Information Figure S2 - PTBP1 is necessary for optimal activation of CD8 T cells** A) Cell Trace Violet profiles (left) of control (Live eFluor780$^-$, CD8a$^+$ cells from a *CD4$^{CreTg}$* mouse) and PTBP1-deficient (Live eFluor780$^-$, CD8a$^+$ cells from a *CD4$^{CreTg}$ Ptbp1$^{fl/fl}$* mouse) naïve cells cultured *ex vivo* with plate-bound anti-CD3 (5 µg/ml) and anti-CD28 (1 µg/ml) antibodies for 48h. Numbers of cells in each number of divisions are shown on the right. Data shown is from one representative experiment out of four. Shown are padj values from a Sidak's multiple correction test post a repeated measures two-way ANOVA. B) IL-2 amounts detected in cultures of naïve CD8 T cells from mice with the indicated genotypes after 24h stimulation on plates pre-coated with 5 µg/ml anti-CD3 with 1 µg/ml anti-CD28 antibodies. Each point shows data from supernatants of cells isolated from one mouse. C) Cell Trace Violet profiles (left) of control (Live eFluor780$^-$, CD8a$^+$ cells from a *CD4$^{CreTg}$* mouse) and PTBP1-deficient (Live eFluor780$^-$, CD8a$^+$ cells from a *CD4$^{CreTg}$ Ptbp1$^{fl/fl}$* mouse) naïve cells cultured *ex vivo* with plate-bound anti-CD3 (5 µg/ml) and anti-CD28 (1 µg/ml) antibodies in the presence of exogenous IL-2 (20 ng/ml) for 48h. Numbers of cells in each number of divisions are shown on the right. Data shown is from one representative experiment out of four. Shown are padj values from a Sidak's multiple correction test post a repeated measures two-way ANOVA. D) Graphs showing Proliferation Indexes and % of Divided from the histograms on **Fig S2A** and **S2C**. Each point shows data from CD8 T cells isolated from one mouse. Data shown is from one experiment. E) Percentages of active-Caspase-3+ cell identified by flow cytometry amongst naïve CD8 T cells labelled with Violet Cell Trace and stimulated *ex vivo* with plate bound anti-CD3 (5 µg/ml) and anti-CD28 (1 µg/ml) with exogenous IL-2 (20 ng/ml) for the indicated time points. Each point shows data from an individual mouse. Bars show arithmetic means. FDR-adjusted p values from a Sidak's multiple correction test post a repeated measures two-way ANOVA are shown. Data shown are from two independent experiments.

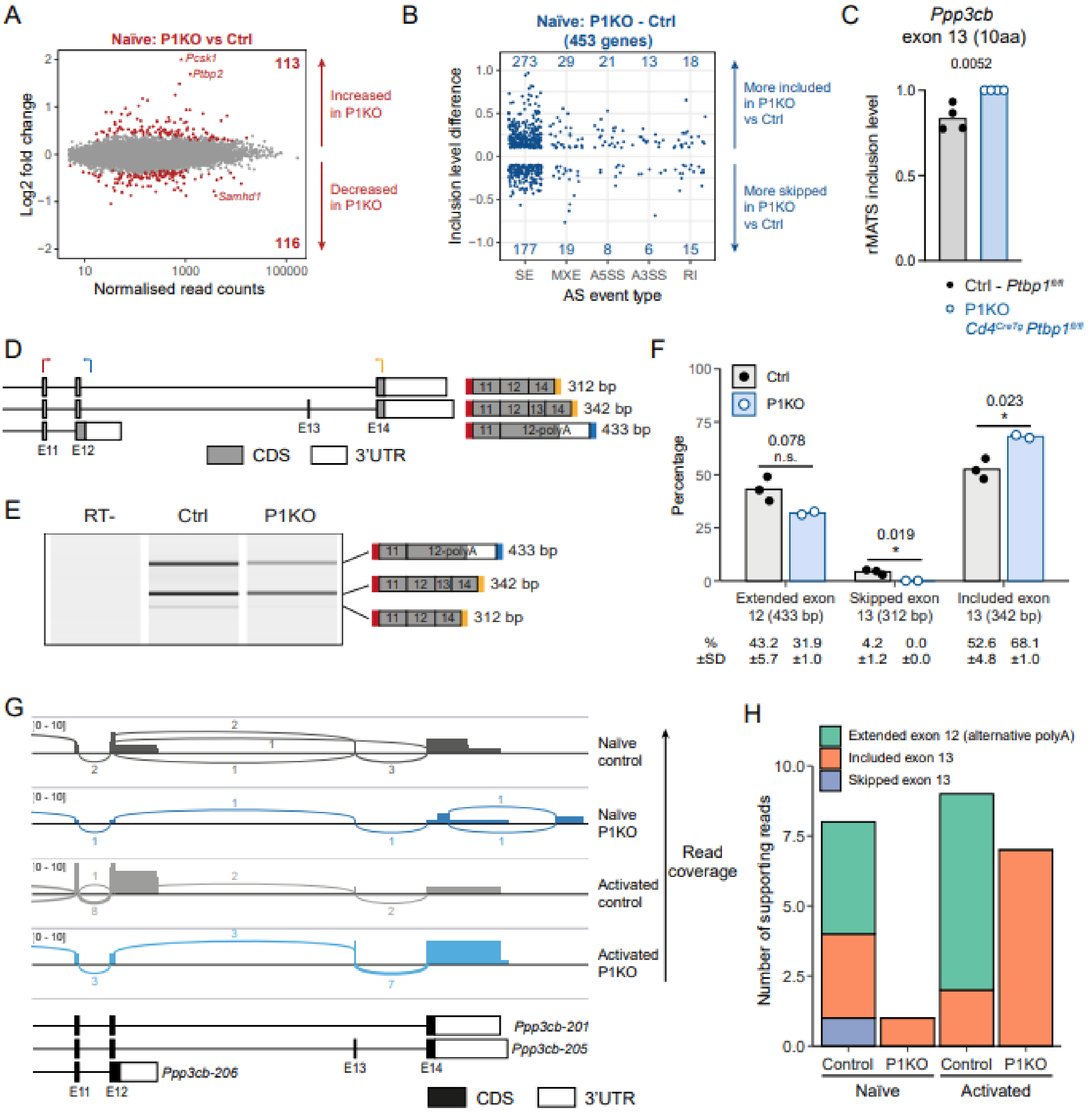


**Supporting Information Figure S3 - PTBP1 regulated gene expression programs in CD8 T cells** A) Changes in mRNA abundance due to *Ptbp1* deletion calculated with DESeq2. Each point shows Log2 FC of an individual gene. Red points show genes with an FDR-adjusted p value <0.05. B) Changes in AS in naïve CD8 T cells due to *Ptbp1* deletion analysed with rMATS. Each point shows the inclusion level differences for an individual event that has changes in AS (|inclusion level difference| > 0.1 and FDR<0.05). Inclusion level differences are grouped by the type of alternative splicing event: skipped exons (SE), mutually exclusive exons (MXE), alternative 5' or 3' splice sites (A5SS and A3SS, respectively) or retained introns (RI). C) rMATS inclusion level of exon 13 in *Ppp3cb* transcript in naïve CD8 T cells from control ($Ptbp1^{fl/fl}$) and P1KO mice ($Ptbp1^{fl/fl}$ $Cd4^{CreTg}$). D) Schematic showing locations of primers used in RT-PCR to amplify *Ppp3cb* transcripts, and the products and sizes expected from different isoforms. E) Virtual gel output from QIAxcel analysis of RT-PCR products amplified from naïve CD8 T cell RNA using the primers shown in D. An RT- control is shown, together with representative samples from a control (n = 3) and P1KO (n = 2) mouse. F) QIAxcel quantitation of the percentage detected of the three RT-PCR products amplified from different *Ppp3cb* isoforms, as shown in D-E. G) Sashimi plot of *Ppp3cb* transcripts in naïve and activated CD8 T cells from 4 (naïve) or 3 (activated) merged replicate datasets for control ($Ptbp1^{fl/fl}$) and P1KO mice ($Ptbp1^{fl/fl}$ $Cd4^{CreTg}$), showing Oxford Nanopore Technologies long read mRNA sequencing coverage of reads that overlap with at least one *Ppp3cb* exon and contain at least one splice junction. Arches and numbers indicate the number of reads mapping to a given junction. H) Summary data for the total number of reads visualised in G that could be unambiguously assigned to *Ppp3cb* isoforms that included or skipped exon 13, or that utilised the alternative polyadenylation site within an extended exon 12.

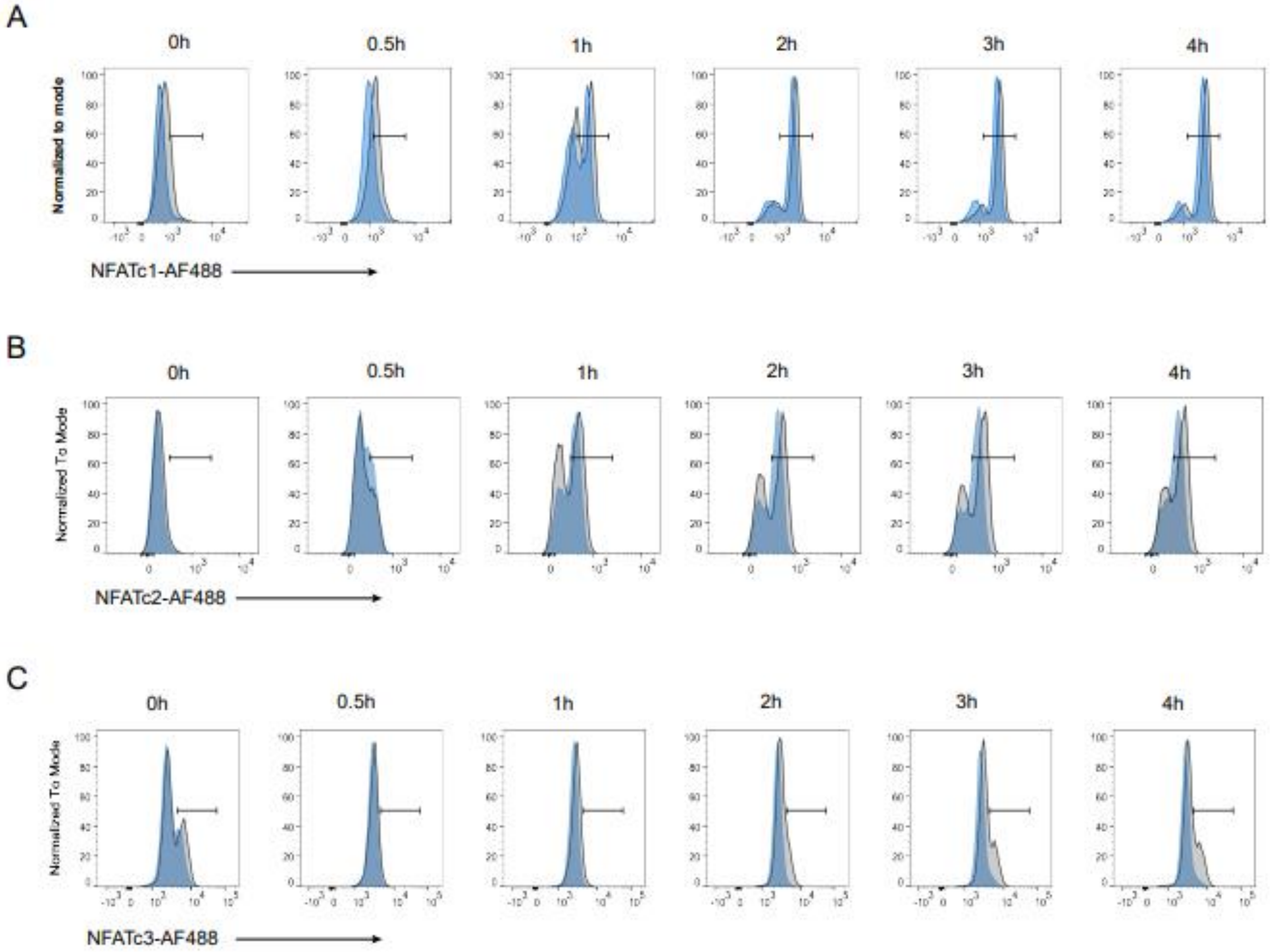


**Supporting Information Figure S4 - PTBP1 regulates nuclear translocation of NFAT** Flow cytometry staining of NFATc1 (A), NFATc2 (B) and NFATc3 (C) in isolated nuclei from control or P1KO mouse naïve CD8 T cells cocultured with bulk splenocytes pulsed with 10 nM of OVA peptide at the time indicated. Gates show positive populations. OT-I nuclei were identified as CellTrace Violet$^{hi}$ CD3e$^{low}$.
